# Assessing extraction methods and diversity of anthocyanins from purple-fleshed sweet potatoes grown in cooler climates

**DOI:** 10.1101/2020.08.23.262808

**Authors:** Alexandra A. Bennett, Kai Fan, Gaurav D. Moghe

## Abstract

Anthocyanins are economically valuable phytochemicals of significant relevance to human health. Multiple fruit and vegetable sources for industrial-scale anthocyanin purification exist, however, each source has distinct anthocyanin levels and profiles conferred by modifications to the central anthocyanidin core. In this study, we assessed three purple-fleshed and one orange-fleshed cultivars of sweet potato, with the goal of studying their anthocyanin yield and diversity when this warm-weather crop is grown in cooler, northern latitudes. Comparison of multiple anthocyanin extraction methods revealed acidified ethanol extraction of lyophilized roots as the optimal method, producing a high, average yield of ∼800 mg anthocyanins/100g dry weight. Mass spectrometric analysis of sweet potato extracts identified eighteen high-confidence anthocyanins – all derived from peonidin and cyanidin cores – contributing to over 90% of the total anthocyanin signal. The concentrations of different anthocyanins were variable between the three purple-fleshed cultivars, while low anthocyanin accumulation was observed in the orange-fleshed cultivar. Further assessment of the untargeted high-resolution mass spectrometry data using MS/MS molecular networking revealed existence of low-abundance anthocyanins with delphinidin and pelargonidin cores, as well as over 250 peaks comprising of potential anthocyanins and flavonoids. These results provide a comprehensive insight into anthocyanin yields of purple-fleshed sweet potato grown in the northern latitudes and reveal the large diversity of anthocyanins and flavonoids in this popular crop.

## Introduction

Anthocyanins are water-soluble phytochemical pigments of significant health and economic value, which belong to a class of polyphenolic compounds called flavonoids. Found in many fruits and vegetables, flavonoids possess antioxidant activities of benefit in managing ageing, stress, cancer and other health conditions which makes them desirable for cosmetic, nutritional and health industry applications^1–3^. Because of their color properties, anthocyanins are also of significant interest as natural food coloring agents^3,4^. Due to these diverse uses, the global market for flavonoids is expected to exceed $1 billion by 2026^5^.

Flavonoids are downstream products of the phenylpropanoid pathway and are defined by presence of a flavone ring structure comprised of three rings termed A, B and C^6^. Anthocyanidins – the aglycone of anthocyanins – are differentiated from other flavonoids by the presence of a positively charged oxygen on the C ring which results in a flavylium cation. The positive charge of the ring confers color and chemical instability to anthocyanidins. Modification through glycosylation, acylation, methylation, and hydroxylation reactions can stabilize these compounds, making them more resistant to changes in pH, temperature and ultraviolet light, and thus, more useful for commercial applications^4,6^. The most common modification is glycosylation, producing anthocyanins. Currently, the major sources of industrial production of anthocyanins include skins of grapes processed in the wine industry, berries, black carrots, red cabbage, and purple-flesh sweet potatoes^7,8^.

Sweet potato (*Ipomoea batatas*) is an attractive crop for anthocyanin extraction. While the major cultivated varieties are orange-fleshed and rich in carotenoids such as β-carotene^9^, there are also purple-fleshed varieties. One meta-analysis compared anthocyanin yields from various sources and estimated 84-174 mg anthocyanins/100 g fresh weight from purple-flesh sweet potatoes compared to 6-600 mg/100 g fresh weight from grapes and ∼25 mg/100 g fresh weight from red cabbage^10^. However, unlike fruits, sweet potato roots produce high biomass, store better, and can be cultivated on large scale. In addition, they primarily make more stable, acylated anthocyanins^10,11^, unlike fruit anthocyanins, which are primarily non-acylated^4,12^. At least 27 anthocyanins are previously known to be present in different sweet potato cultivars^11,13–15^, in addition to other health promoting phenolic compounds^13^.

Multiple studies have previously characterized extraction processes and anthocyanin content from different sweet potato varieties. Processing methods utilized include homogenizing raw tissue, freezing, lyophilization, boiling, steaming, roasting and frying^16^. After processing, anthocyanins can be extracted using organic solvents^17^, alkaline solution, resins^18^, accelerated solvent extraction^14^ and sonication^19^, among other methods. Ethanol, methanol, and acetone:chloroform have been tested as solvents for anthocyanin extraction, though ethanol is generally preferred for food-grade anthocyanin purification because of methanol toxicity^20^. Studies in sweet potato have revealed primarily peonidin and cyanidin derived anthocyanins^14,16^, but the overall extent of anthocyanin diversity is not known.

In the United States, sweet potato is a crop primarily grown in warmer states such as North Carolina – which leads the country in sweet potato production^21^ – Louisiana, Mississippi, and California, also because of the longer growing season. These states accounted for >90% of the 3.1 billion pounds of sweet potato grown in 2015 in the USA^21^. Originating from tropical areas in Central America and northwestern South America^22^, sweet potato is adapted to warmer climates and soils and a longer growing season. However, many regions in the northern latitudes in the US and Canada are increasingly interested in cultivation of this crop due to its economic value in grocery markets as well as for industrial products^23–26^. Geneva, New York, where this project was conducted has, on an average, ∼5 °C lower day time temperatures and 5-12 °C lower night time temperatures in the primary growing months of June-September, than principal purple sweet potato growing regions such as Raleigh, North Carolina (USA), Xuzhou, Jiangsu (China) and Naha, Okinawa (Japan) **(Supplementary Fig. 1)**. This temperature gap is representative of such a gap across the northern latitudes, thus necessitating the evaluation of sweet potato growth and anthocyanin yield in cooler climates for a better understanding of their economic potential.

Thus, in this study, we focused on the anthocyanin yield and content of one orange-fleshed and three purple-fleshed varieties grown in the cooler climate of upstate New York. One advantage of growing sweet potatoes in the north is a relative absence of natural pests, which leads to less pesticide application and a resulting organic cultivation. Furthermore, cold stress is known to induce anthocyanin production in other species^27–29^. While high yield orange-fleshed varieties such as Beauregard and Covington have been studied in more detail for their growth characteristics^30–32^, the anthocyanin content of purple-flesh varieties grown in the north has not been assessed. Thus, the goals of this study were to establish an anthocyanin extraction method, estimate overall anthocyanin yield, and identify the different types of anthocyanins present in three purple-fleshed sweet potato varieties grown in upstate New York. We used a combination of spectrophotometric, mass spectrometric and computational methods to understand the amounts and types of anthocyanins and flavonoids in these varieties. Our results suggest substantial diversity and yield of these compounds, reinforcing the value of purple sweet potatoes as industrial sources of anthocyanins and flavonoids.

## Materials and Methods

### Growing conditions

*Ipomoea batatas* (sweet potato) slips of four cultivars were obtained as follows: ‘Kotobuki’ and ‘Purple Passion’ from George’s Plant Farm, and ‘All Purple Sweet Potato’ and ‘Beauregard’ from Southern Exposure Seed Exchange. Slips were transplanted to “Cornell Mix” soil substrate and maintained within a greenhouse at Cornell University until field season, under summer conditions. Slips were transplanted to a field in Geneva, NY onto raised plastic beds with 1.8 m centers and 45 cm spacing. Plants were maintained with drip irrigation and fertigation until harvest after 106 days. Sweet potato roots were cured for 3 weeks at room temperature, washed and were kept at ∼60-65 °C for another 6 weeks. Sweet potatoes were finally moved to long term storage at 55 °C at ∼80% relative humidity. The weight, length, and circumference were measured and recorded one month after harvest for each sweet potato.

### Solvents and chemicals

ACS grade reagents ≥99.8% methanol, 95% ethanol, ≥88% formic acid and ≥99.7% glacial acetic acid, 36.5-38% hydrochloric acid, potassium chloride as well as ≥99% sodium acetate were sourced from VWR, Radnor, Pennsylvania, USA. The standard ≥90% cyanin chloride was purchased from Santa Cruz Biotechnology, Dallas, Texas, USA. Ultra-high purity water was generated by an Elga PureLab Ultra reverse osmosis system equipped with a LC182 purification cartridge.

### Standardization of solvent extraction

First, 1.5 kg of *I. batatas* cultivar ‘Kotobuki’ was selected from storage at random at ∼40 days after harvest. Sweet potatoes were divided into five replicates of approximately 300 g each. Samples were cut into approximately 5 mm thick discs, wrapped in tin foil to minimize anthocyanin leakage in water, and boiled at 110 °C for 40 minutes (time beginning after water returned to a boil). After cooking, potatoes were homogenized in a Waring 7011G blender on high, snap frozen in liquid nitrogen, and stored in a −80 °C freezer. Frozen tissue was then ground to a powder with a Victoria cast iron grain mill that had been chilled with liquid nitrogen. Samples were mixed with dry ice to keep tissue frozen. Ground samples were kept in the −80 °C freezer over a span of two days to allow dry ice to completely vent off. Frozen powder was divided into two 100 g portions. One portion was extracted in 100 mL 75% methanol with 10% acetic acid while the second portion was extracted in 100 mL 75% ethanol with 10% acetic acid.

### Cooking methods comparisons

Cultivar ‘All Purple Sweet Potato’ was weighed into five replicates of approximately 350 g each and sliced into approximately 5 mm thick discs. These slices were then mixed randomly and split into three 100 g portions. Samples were wrapped in tinfoil and labeled. One portion from each replicate was then boiled at 110 °C for 40 minutes. A second portion was boiled at 110 °C for 20 minutes and then submerged in ice cold water for 20 minutes. The last portion was pressure cooked in water with a Fagor LUX Multi-Cooker (670041880) for 20 minutes (time beginning after pressure and temperature was reached). All samples were then drained of excess water, placed in labeled Ziplock bags, and homogenized by hand within the bags. Samples were flattened into thin sheets and frozen in a −80 °C freezer. These frozen samples were then fine ground in a grain mill with the aid of dry ice. Dry ice was allowed two days to vent off completely in the −80 °C freezer. Samples were extracted in 100 mL of 75% ethanol with 10% acetic acid.

### Overall processing methods comparisons

Five randomized replicates from sweet potato cultivar ‘All Purple Sweet Potato’ were sliced into 5 mm thick discs and mixed. Each replicate was divided into four portions of ∼100 g each for four different processing methods (raw, frozen, lyophilized, and pressure cooked). The raw portion was ground in a grain mill and extracted. The second and third portions were snap frozen in liquid nitrogen. The second portion was then ground with the aid of dry ice and given two days to vent before extraction. The third portion was lyophilized over a 24-hour period and weighed before grinding. The last portion was processed by pressure cooking as previously described and then weighed. 50 g of each method was weighed out (with lyophilized and pressure-cooked weights adjusted to reflect fresh weight). Samples were extracted in 50 mL of 75% ethanol with 10% acetic acid.

### Cultivar comparisons

Four replicates of the four cultivars under study were processed via lyophilization method as described above, with the exception that 20 g dry weight was extracted in 50 mL.

### Anthocyanin extraction

Extraction was done in dark. Homogenized tissue was extracted in designated solvent for 60 minutes on an VWR® Variable Speed Rocker set to the highest speed. Liquid was vacuum filtered with a Buchner funnel lined with Whatman’s student grade filter paper and stored at −20°C overnight until anthocyanins were quantified with a spectrophotometer.

### Quantification of anthocyanins

Cyanin chloride (referred to here as cyanin) was dissolved in methanol containing 1% formic acid to a concentration of 8 mM. 40 μL of cyanin was then mixed with 1960 μL of 25 mM potassium chloride pH 1.0, and another 40 μL cyanin was mixed with 1960 μL of 0.4M sodium acetate pH 4.5. Cyanin was given 15 minutes to reach equilibrium. Cyanin was then diluted two-fold serially to generate 7 concentrations ranging from 160 μM to 1.25 μM (this reflects a sample concentration of 8 mM to 62.5 μM). Cyanin solutions were then measured in triplicate with a Varian Cary 50 Bio UV-visible spectrophotometer within 1 hour and analyzed via pH differential method^33^. The computer system monitored and analyzed data using Varian Cary WinUV Simple Reads software version 4.10 (build 464). The same protocol was used for measuring anthocyanins from root extracts. Depending on the pH, potassium chloride or sodium acetate was used as blanks. Equilibrated sample and standard curve absorbance measurements were then taken at 520 nm and 700 nm by the spectrophotometer^33^. Final absorbance values were calculated through the pH differential method as follows:

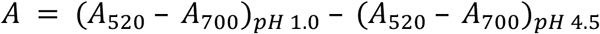

Kolmogorov-Smirnov test was performed in Statistical Analysis System’s (SAS) JMP Pro software v14.3.0. Cyanin values were used to generate a standard curve. The linear regression from this standard curve was used to calculate mg/g concentrations of each condition and cultivar in Microsoft Excel 365.

### Estimation of concentrations and dilution corrections

For processing comparisons, samples were reported in fresh weight equivalence. To make the experiment comparable across all four conditions, dilution for pressure cooking and concentration for lyophilization needed to be accounted for. This was done through dilution equations:

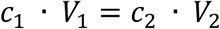

In this equation, *c*_1_ is the original concentration of the raw extract measured and *V*_1_ is the original volume of the extract accounting for the 50 mL of solvent used and (if applicable) the water in the tissue and the water gained by cooking. *c*_2_, what is being solved for, reflects what the concentration of the sample would be if it was in the 50 mL of solvent plus the water contained within the tissue as this is the amount of solution a fresh weight extract would be in. Thus, *V*_2_is the volume made up of 50 mL of solvent and the amount of water in the tissue. The amount of water in the pressure-cooked sample was calculated by taking the average percent water (43%) measured in lyophilized samples. Calculations were done this way to maintain comparability to other fresh weight studies.

### Mass spectrometric analysis

One mL of anthocyanin extract was transferred to amber HPLC vials (VWR 46610-726). Samples were separated with a Dionex UltiMate 3000 UHPLC system and Phenomenex Kinetex F5 column (00D-4722-AN, 1.7µm particle size, 100Å pore size, 100 mm length, 2.10 mm ID) at a flow rate of 0.6 mL/min. Solvent A was UHP H_2_O and Solvent B was acetonitrile, both with 3% formic acid. Solvent gradient was as follows (values in Time[minutes]: %B): 0.0: 5%, 1.0: 12%, 7.5: 15%, 8.0: 40%, 9.0: 14%, 9.0: 5%, 10.0: 5%. After separation, anthocyanins were detected by a Dionex UltiMate 3000 diode array and multiple-wavelength detector (DAD) at 520 nm with a 700 nm reference in addition to US-VIS full spectrum. A Thermo Fisher Orbitrap QE detected anthocyanins through data dependent MS2 (DDMS2) scans after the DAD. 520 nm is a midrange maximum absorbance (λ_max_) value amongst anthocyanins while 700 nm is used to correct for haze as there is no absorbance by anthocyanins at this wavelength

Raw data was converted to Analysis Based File (ABF) format using Reifycs ABF Converter and imported into MS-DIALv4.24^34^. Data from samples was filtered and aligned using MS-DIAL and data on all aligned metabolites was exported in mascot generic format (MGF). Anthocyanins were selected manually in MS-DIAL based upon high abundance aglycone signature fragments (*m/z*: 287.06 for cyanidin, 271.06 for pelargonidin, 303.05 for delphinidin, 301.07 for peonidin and 317.07 for petunidin) and exported to MS-Finder version 3.44 to obtain their chemical formulas. ThermoFisher Chromeleon was used to manually align 520 nm peak data from the DAD to a subset of high intensity MS anthocyanin peaks previously identified. Peak intensities from UV-VIS data was used to calculate percent anthocyanin composition in Chromeleon.

### Molecular networking analysis

All metabolites exported in MGF format from MS-DIAL were filtered using a custom Python script to identify anthocyanin-like peaks (including flavonoids of similar masses) across all lyophilized cultivar samples. This script selected all peaks with both the anthocyanidin and its mono or disaccharide derivative as peaks with intensity>3000. For anthocyanidins, we used the *m/z* values for all cores mentioned above as well as malvidin (*m/z*: 331) and rosinidin (*m/z*: 315). Monosaccharides considered included all masses for hexose, deoxy hexose and pentose sugars, while disaccharides were pairwise combinations of the monosaccharide masses. The filtered MGF file from this script was imported into MS-Finder and the previously hand identified anthocyanins were confirmed to be in this import. MS-Finder was used to generate a molecular network (MS/MS similarity cutoff of 70%) and export the nodes and edges. Nodes and edges were imported into Cytoscape v3.8.0^35^ for figure generation.

## Results

### Biometric assessment and sweet potato yield

Three purple-fleshed and one orange-fleshed sweet potato varieties were grown from slips in Geneva, NY for 106 days, and were phenotyped after harvesting **(Figure 1; Supplementary File 1)**. Yield metrics for ‘All Purple Sweet Potato’ and ‘Purple Passion’ were not significantly different (Kolmogorov-Smirnov [KS] test, p=0.84), however, ‘Kotobuki’ performed better than these two varieties **(Figure 1A)**. While it had a similar average root weight, circumference, and length, the average number of sweet potatoes per plant was 9.4 – compared to 7.3 for the other two – resulting in greater yield per plant. The orange-fleshed cultivar ‘Beauregard’ outperformed all three purple varieties. The average Beauregard plant produced sweet potatoes that were almost 2X as heavy and 1.5X as wide as ‘Kotobuki’, despite having the lowest average number of sweet potatoes harvested per plant (4.6). The sweet potatoes of the four varieties were used to identify an optimal anthocyanin extraction method and assay their anthocyanin profiles.

**Figure 1:**
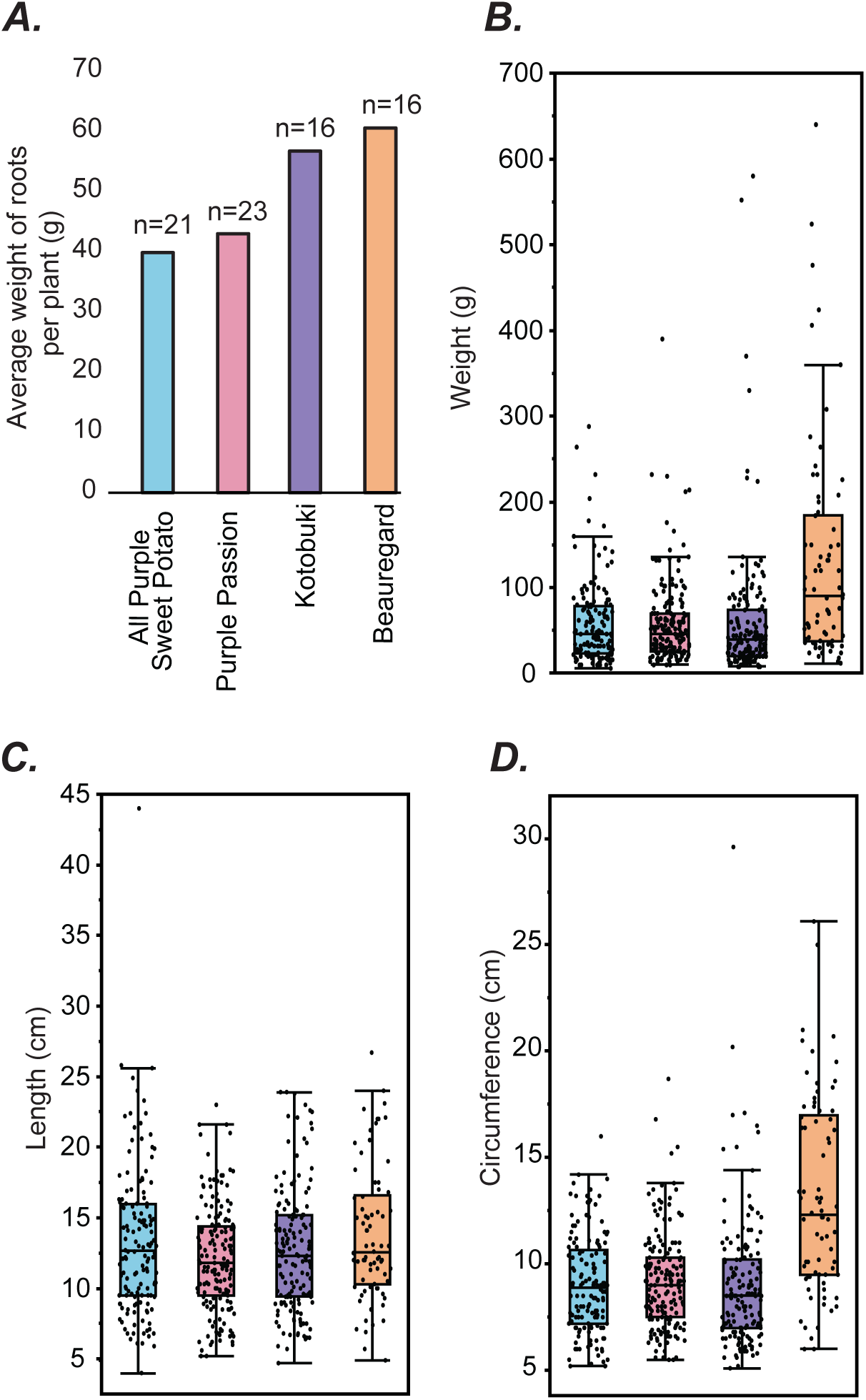
Phenotyping of studied sweet potato cultivars. (A) Average grams roots per plant, obtained by dividing total weight of all sweet potatoes for a given cultivar by the number of plants harvested. Number of plants for each cultivar is indicated above the bar graph. (B-D) Weight, length and circumference distributions for each sweet potato variety. Colors used for each cultivar are the same across all panels.

### Optimization of anthocyanin extraction process

Anthocyanins are relatively hydrophobic and have poor extraction abilities in a neutral pH aqueous solution. Thus, solvents such as acidified ethanol, acidified methanol, acetone:chloroform, are typically used in the extraction of anthocyanins^10^. Given material cost is an important consideration for future industrial applications, we focused on only the first two solvents, and compared the ability of 75% acidified ethanol versus 75% acidified methanol to extract anthocyanins from boiled ‘Kotobuki’ sweet potatoes at room temperature, using cyanidin-3,5-diglucoside as an external standard **(Figure 2A; Supplementary File 2)**. Ethanol (median 1.65 mg/g fresh weight) was not significantly better at extracting anthocyanins than methanol (median 1.55 mg/g fresh weight) **(Figure 2B)** (KS test, p=0.3571). This result is in agreement with a previous study that compared anthocyanin concentrations in alcoholic sweet potatoes extracts at 25 °C and found no significant difference^10^. Our results suggest that prior temperature processing has no differential effect on the abilities of these solvents to extract total anthocyanins.

**Figure 2:**
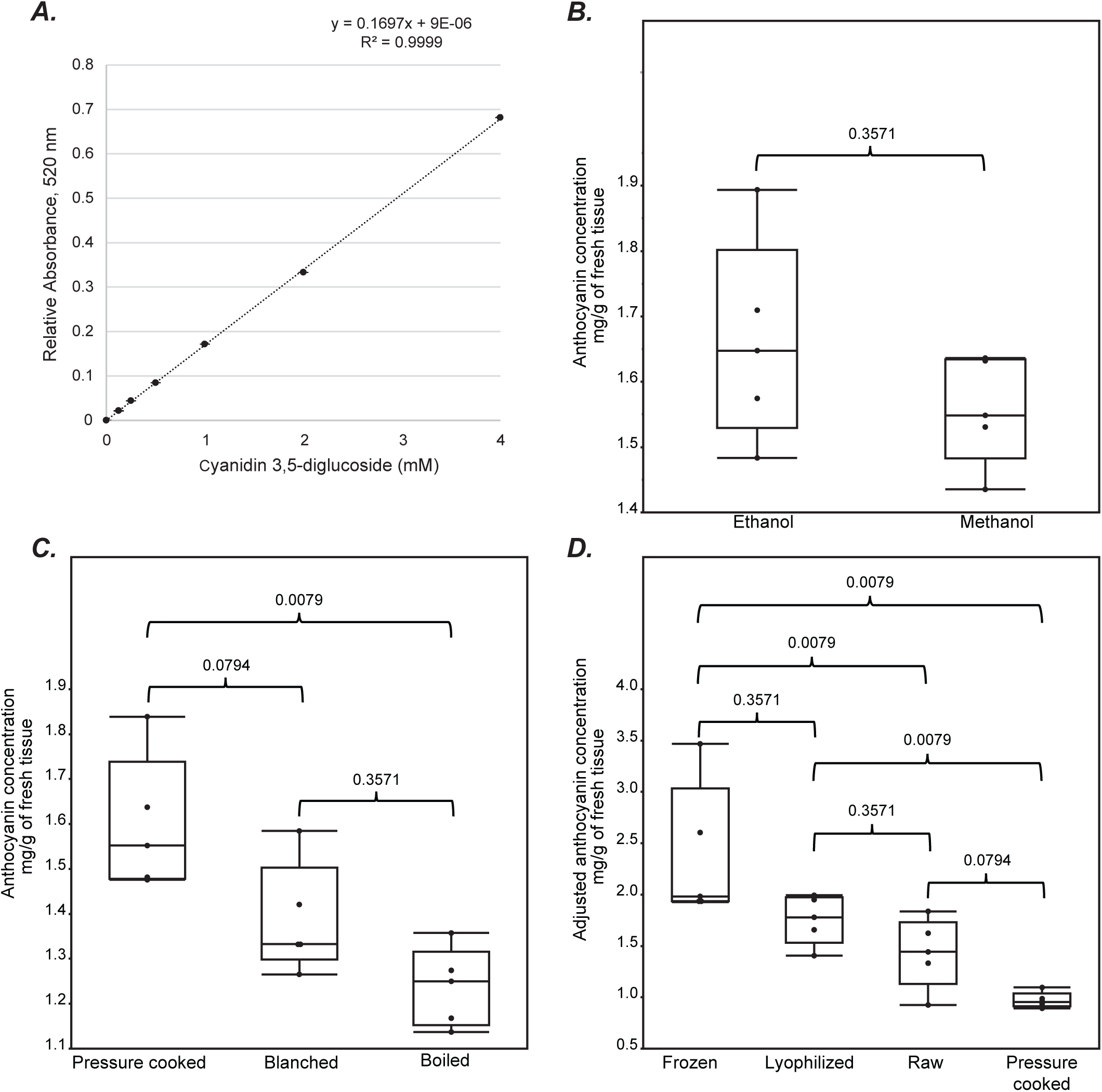
Quantification of different anthocyanin processing methods. using (A) a standard curve of representative anthocyanin cyanidin 3,5-diglucoside (cyanin). Total monomeric anthocyanin yields (cyanin equivalence) of purple fleshed sweet potatoes were compared among three extraction and processing experiments. These were (B) 75% ethanol versus 75% methanol (both containing 10% acetic acid) extraction of cultivar ‘Kotobuki,’ (C) different processing methods of boiling, blanching, and pressure cooking of ‘All Purple Sweet Potato’, and (D) freezing, lyophilizing, pressure cooking, and raw of ‘All Purple Sweet Potato.’ Adjustments performed noted in the Methods section. Non-parametric Kolmogorov-Smirnov test was utilized to assess p-values.

Using acidified 75% ethanol as the extraction solvent – selected due to significantly lower toxicity than methanol – we next compared the impact of different cooking methods, namely boiling, blanching and pressure cooking, on anthocyanin levels from sweet potatoes. Previous studies suggest mixed results, with either raw extraction or boiling as the optimal method for anthocyanin release from cells^36–39^. The cooking process denatures polyphenol oxidase enzymes and disintegrates cell and vacuole membranes to release anthocyanins. All three cooking methods yielded significantly different results from one another in cultivar ‘All Purple Sweet Potato’, with pressure cooking resulting the highest total monomeric anthocyanin median yield (1.55 mg/g fresh weight) **(Figure 2C)**. Blanching and boiling yielded, ∼80-85% of total anthocyanins vs. pressure cooking. While pressure cooking was the best performing cooking method, previous studies have been conflicting on if these high temperatures have a positive impact on total intact sweet potato anthocyanin content of the final extract or not^37,38,40^. Hence, we separately compared pressure cooking with other processing methods that do not require heating – namely homogenization of raw tissue, snap freezing, and lyophilization – at the same post-harvest stage. Freezing and lyophilization are both low temperature methods that may not only preserve anthocyanin levels but also aid in anthocyanin release due to cellular rupture.

Pressure cooked samples yielded a median 0.96 mg/g fresh weight. The lower yield of this batch of pressure cooked sweet potatoes compared to the previous experiment (1.55 mg/g) could likely be an effect of anthocyanin degradation due to additional two weeks of sweet potato storage between the two experiments (2 months vs. 2.5 months), potentially resulting in degradation^41^. In this analysis, pressure cooking resulted in lower anthocyanin yield than sweet potatoes that were extracted raw with no freezing step (1.44 median mg/g), which itself was lower than both frozen and lyophilized methods **(Figure 2D)**. Freezing samples in liquid nitrogen with (1.78 median mg/g fresh weight) or without (1.98 median mg/g fresh weight) subsequent lyophilization resulted in the highest levels of anthocyanins extracted. Although raw frozen had slightly higher yield than lyophilized, the standard deviation for freezing without lyophilization (0.67 mg/g) was >2X freezing with subsequent lyophilization (0.24 mg/g). This suggested that lyophilization resulted in more reproducible and high anthocyanin yields, possibly because of the variable water content in samples without lyophilization. Thus, sweet potatoes for further experiments were snap frozen and lyophilized before extraction in 75% acidified ethanol. After standardizing this optimal anthocyanin processing method in one variety, we used this to assess the anthocyanin variability among different sweet potato cultivars.

### Quantification of monomeric anthocyanin content in purple sweet potato cultivars

While relative anthocyanin concentration diversity and phenotyping among sweet potatoes is visually apparent, absolute concentrations are not. The orange cultivar, ‘Beauregard,’ contained no accurately quantifiable levels of anthocyanins neither visually nor with a spectrophotometer. The three purple cultivars analyzed, ‘Kotobuki’ (7.37 median mg/g dry weight), ‘All Purple Sweet Potato’ (7.25 median mg/g dry weight), and ‘Purple Passion’ (8.23 median mg/g dry weight), contained levels of anthocyanins that were not significantly different from one another **(Figure 3A; Supplementary File 3)**, suggesting relative uniformity in the processes that lead to anthocyanin accumulation. While the overall anthocyanin levels were same, it is possible that the specific anthocyanin content is different in each cultivar. Thus, we used LC-MS to further characterize the types of anthocyanins present in these sweet potatoes.

**Figure 3:**
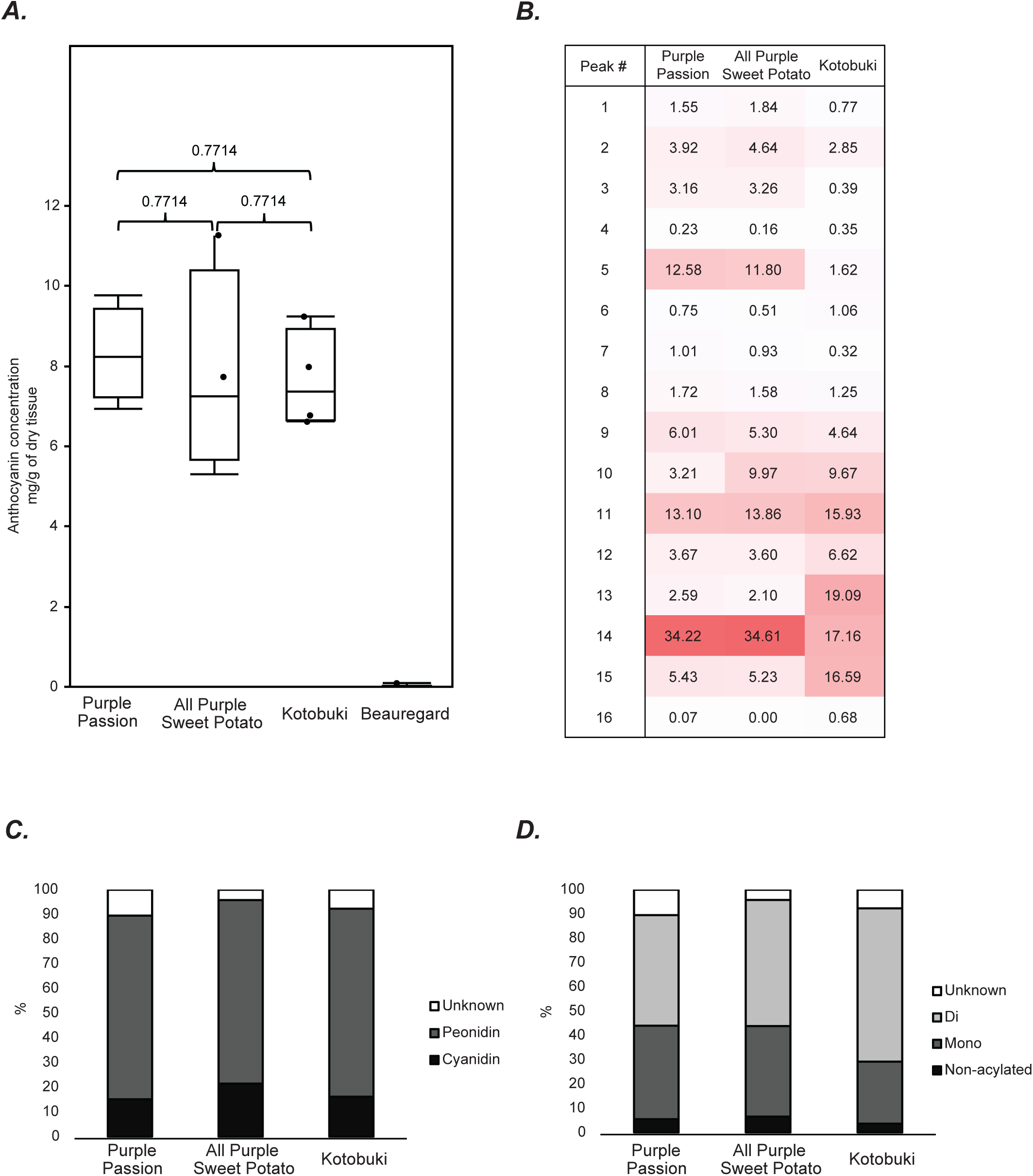
Anthocyanin diversity among the four assessed cultivars. (A) Box plot of total monomeric anthocyanin concentrations (cyanin equivalence) of sweet potato cultivars ‘Kotobuki,’ ‘All Purple Sweet Potato,’ ‘Purple Passion,’ and ‘Beauregard.’ KS test was utilized to assess p-values. (B) Heatmap showing percent of total anthocyanin content for each identified anthocyanin, based on UV-VIS 520 nm data. (C) Peonidin and cyanidin are the major aglycone classes found in the studied sweet potatoes (D) the majority of anthocyanins measured were acylated to some degree.

### Analytical characterization of anthocyanins and peak identification

Anthocyanins were identified from whole root extracts using a combination of spectrophotometric and mass spectrometric methods **(Figure 4A,C)**. The association of absorption spectrometry data with LC-MS/MS fragmentation data led to identification of sixteen high-confidence anthocyanin peaks representing eighteen anthocyanins **(Table 1)**. Sweet potatoes are known to produce acylated anthocyanins, which are chemically more stable to environmental changes than non-acylated anthocyanins. Acylation of anthocyanins can be determined using the ratio between the λ_max_ peak (∼520 nm) and the acylation peak (∼330 nm), as acylation results in increased absorptivity (hyperchromic effect) of the acylation peak^12^. An example of this phenomenon is shown for cyanidin 3-caffeoyl-p-hydroxybenzoyl sophoroside-5-glucoside **(Figure 4B)**. Sixteen of the eighteen peaks were found to be acylated.

**Table 1:**
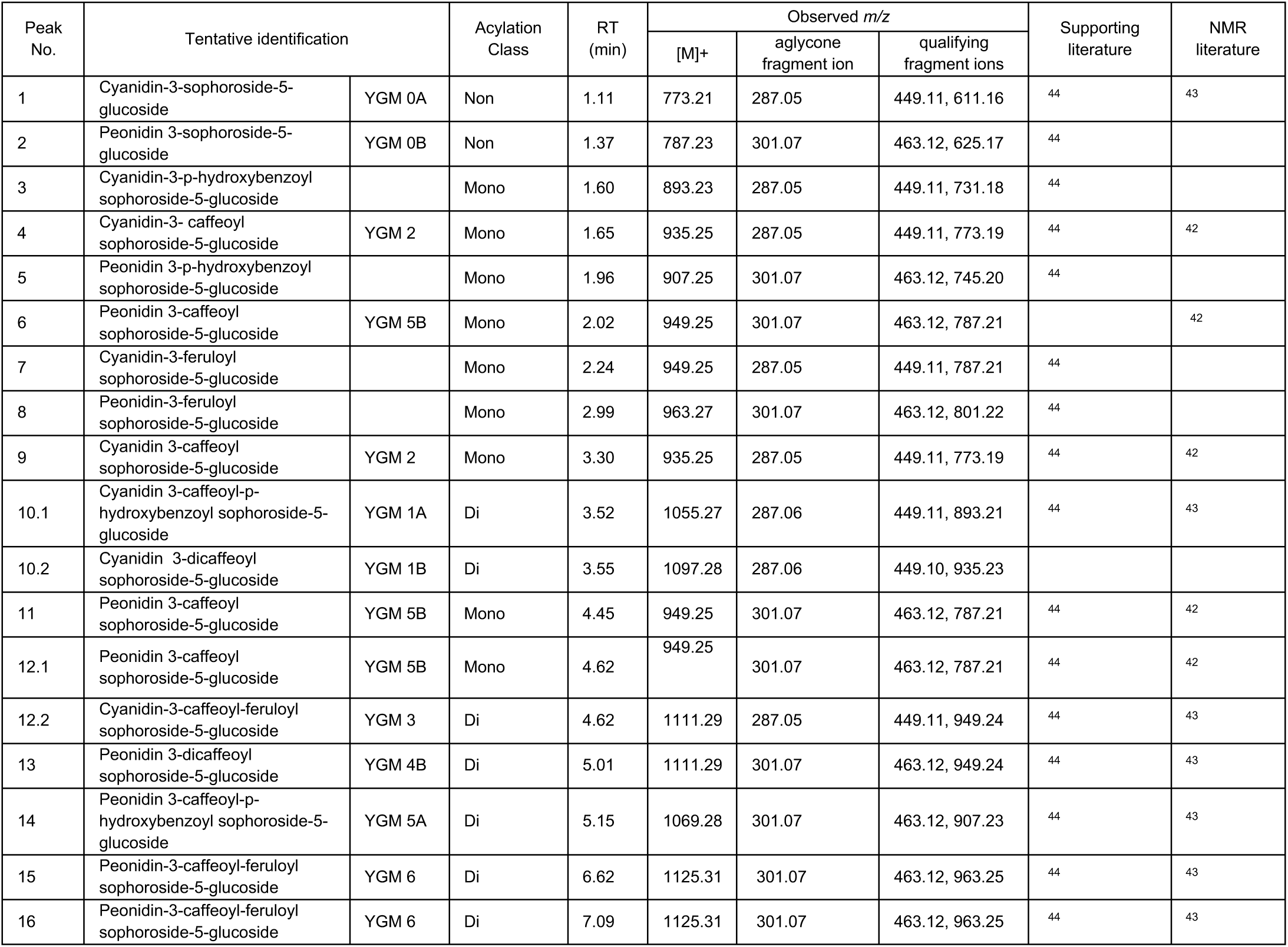
Anthocyanins identified across the three purple sweet potato varieties. YGM stands for ‘Yamagawamurasaki,’ the cultivar from which the previous sweet potato anthocyanins were elucidated from via NMR^43^. Retention time is abbreviated to RT.

| Peak No. | Tentative identification |  | Acylation Class | RT (min) | Observed m/z |  |  | Supporting literature | NMR literature |
| --- | --- | --- | --- | --- | --- | --- | --- | --- | --- |
|  |  |  |  |  | [M]<sup>+</sup> | aglycone fragment ion | qualifying fragment ions |  |  |
| 1 | Cyanidin-3-sophoroside-5-glucoside | YGM 0A | Non | 1.11 | 773.21 | 287.05 | 449.11, 611.16 | 44 | 43 |
| 2 | Peonidin 3-sophoroside-5-glucoside | YGM 0B | Non | 1.37 | 787.23 | 301.07 | 463.12, 625.17 | 44 |  |
| 3 | Cyanidin-3-p-hydroxybenzoyl sophoroside-5-glucoside |  | Mono | 1.60 | 893.23 | 287.05 | 449.11, 731.18 | 44 |  |
| 4 | Cyanidin-3-caffeoyl sophoroside-5-glucoside | YGM 2 | Mono | 1.65 | 935.25 | 287.05 | 449.11, 773.19 | 44 | 42 |
| 5 | Peonidin 3-p-hydroxybenzoyl sophoroside-5-glucoside |  | Mono | 1.96 | 907.25 | 301.07 | 463.12, 745.20 | 44 |  |
| 6 | Peonidin 3-caffeoyl sophoroside-5-glucoside | YGM 5B | Mono | 2.02 | 949.25 | 301.07 | 463.12, 787.21 |  | 42 |
| 7 | Cyanidin-3-feruloyl sophoroside-5-glucoside |  | Mono | 2.24 | 949.25 | 287.05 | 449.11, 787.21 | 44 |  |
| 8 | Peonidin-3-feruloyl sophoroside-5-glucoside |  | Mono | 2.99 | 963.27 | 301.07 | 463.12, 801.22 | 44 |  |
| 9 | Cyanidin 3-caffeoyl sophoroside-5-glucoside | YGM 2 | Mono | 3.30 | 935.25 | 287.05 | 449.11, 773.19 | 44 | 42 |
| 10.1 | Cyanidin 3-caffeoyl-p-hydroxybenzoyl sophoroside-5-glucoside | YGM 1A | Di | 3.52 | 1055.27 | 287.06 | 449.11, 893.21 | 44 | 43 |
| 10.2 | Cyanidin 3-dicaffeoyl sophoroside-5-glucoside | YGM 1B | Di | 3.55 | 1097.28 | 287.06 | 449.10, 935.23 |  |  |
| 11 | Peonidin 3-caffeoyl sophoroside-5-glucoside | YGM 5B | Mono | 4.45 | 949.25 | 301.07 | 463.12, 787.21 | 44 | 42 |
| 12.1 | Peonidin 3-caffeoyl sophoroside-5-glucoside | YGM 5B | Mono | 4.62 | 949.25 | 301.07 | 463.12, 787.21 | 44 | 42 |
| 12.2 | Cyanidin-3-caffeoyl-feruloyl sophoroside-5-glucoside | YGM 3 | Di | 4.62 | 1111.29 | 287.05 | 449.11, 949.24 | 44 | 43 |
| 13 | Peonidin 3-dicaffeoyl sophoroside-5-glucoside | YGM 4B | Di | 5.01 | 1111.29 | 301.07 | 463.12, 949.24 | 44 | 43 |
| 14 | Peonidin 3-caffeoyl-p-hydroxybenzoyl sophoroside-5-glucoside | YGM 5A | Di | 5.15 | 1069.28 | 301.07 | 463.12, 907.23 | 44 | 43 |
| 15 | Peonidin-3-caffeoyl-feruloyl sophoroside-5-glucoside | YGM 6 | Di | 6.62 | 1125.31 | 301.07 | 463.12, 963.25 | 44 | 43 |
| 16 | Peonidin-3-caffeoyl-feruloyl sophoroside-5-glucoside | YGM 6 | Di | 7.09 | 1125.31 | 301.07 | 463.12, 963.25 | 44 | 43 |

**Figure 4:**
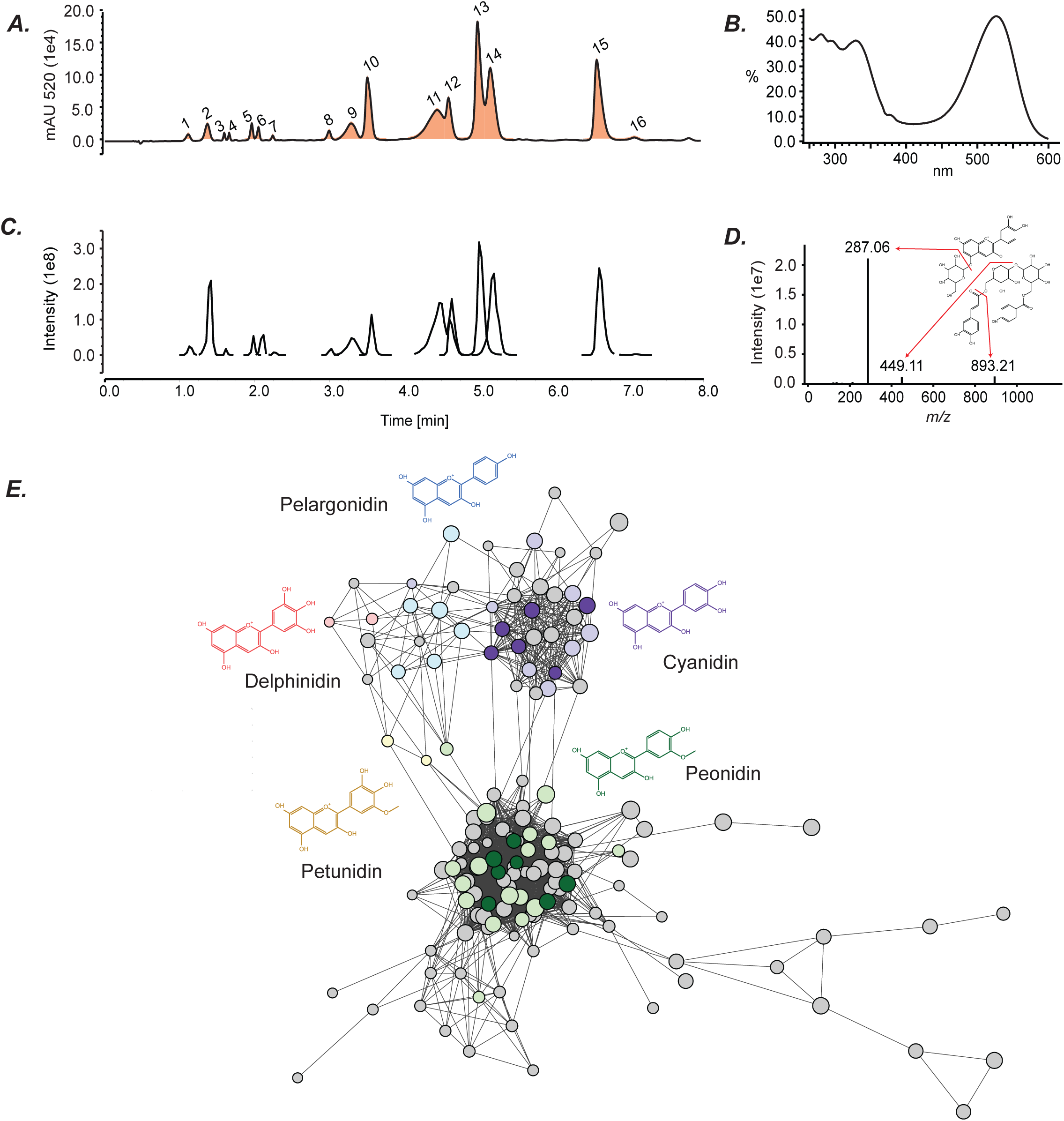
MS/MS analysis of sweet potato anthocyanins. (A-D) Representative LC-DAD-MS data from sweet potato cultivar ‘Kotobuki’ showing (A) a 520 nm UV-Vis chromatogram data is shown with (B) full absorption spectra of of cyanidin 3-caffeoyl-p-hydroxybenzoyl sophoroside-5-glucoside and (C) an extracted ion chromatogram of anthocyanins identified across all sweet potato cultivars with (D) fragmentation pattern of cyanidin 3-caffeoyl-p-hydroxybenzoyl sophoroside-5-glucoside highlighting the cyanidin aglycone major fragment ion of m/z 287.06. (E) MS/MS molecular networking of computationally identified flavonoids. Dark colored nodes represent the 18 high-confidence anthocyanins, and light colored nodes contain similar fragmentation patterns suggesting the same aglycone core containing anthocyanins. See Supplementary Figure 2 for fragmentation patterns. Grey nodes are unidentified (non)-anthocyanin flavonoids. Node size corresponds to number of spectra assimilated within the node.

To identify the anthocyanins, we determined the chemical formula for the eighteen compounds using the parent mass obtained through the high mass accuracy Orbitrap MS. Formulas, retention time, and signature MS/MS fragments – such as the aglycone fragment, mono-glycosylated anthocyanin fragment, acylation patterns – were compared to previous studies^42–44^ for anthocyanin identification. The structure and acylation of the predicted anthocyanins were further confirmed by comparison to NMR studies of known sweet potato anthocyanins. This combinatorial analysis enabled identification of the eighteen compounds **(Table 1)**. Three acyl groups – feruloyl, *p*-hydroxybenzoyl, and caffeoyl – were observed. The high abundance anthocyanins were either peonidin or cyanidin derived and all were 3-sophoroside-5-glucosides with different levels of acylation. Through MS/MS fragmentation, low abundance petunidin, pelargonidin, and delphinidin derived anthocyanins were also detected. While these peaks were too low in abundance to be reliably portrayed in the UV-VIS data, specific signature fragments can still be utilized to tentatively identify low abundance anthocyanins.

### Variation in anthocyanin profiles across purple fleshed sweet potato cultivars

The 18 identified anthocyanins make up >90% of the anthocyanins within our samples across all cultivars sampled. Previous research has shown variation in ratios of these peaks across different distinct sweet potato cultivars^44^. In our analysis, ‘Purple Passion’ and ‘All Purple Sweet Potato’ had very similar compositions with only one anthocyanin, peak 6, being significantly different between the two samples **(Figure 3B)**. In contrast, all but three of ‘Kotobuki’s’ identified anthocyanins (peaks 9-11) were significantly different from anthocyanins in both ‘Purple Passion’ and ‘All Purple Sweet Potato.’

Specific anthocyanins were further assessed for the aglycone type and acylation level. The aglycone type influences the visual pigmentation^45^ while acylation is known to increase stability of anthocyanins due to *pi* stacking interactions between the aromatic rings resulting in intramolecular co-pigmentation^46,47^. The aglycone ratios of all three cultivars assessed were not significantly different, with peonidin making up the largest proportion of the anthocyanin types **(Figure 3C)**. Our results also show that ‘Kotobuki’ had significantly more diacylated anthocyanins and significantly fewer monoacylated anthocyanins when compared to ‘Purple Passion’ and ‘All Purple Sweet Potato.’ All three cultivars have similar levels of non-acylated anthocyanins **(Figure 3D)**.

While anthocyanin levels were below the limit of quantification within the cultivar ‘Beauregard,’ two anthocyanins (peaks 5,14) were above the limit of detection. This indicates low levels of anthocyanin production, possibly occurring due to promiscuous enzymes and/or ‘leaky genes’ within roots of the orange cultivar. All anthocyanins detected in ‘Beauregard’ were peonidin-based and acylated. This result, along with the smaller unknown peaks detected in our LC-MS experiments, indicates the possibility of more low-abundance, unidentified anthocyanins among the purple cultivars.

### Molecular networking of identified and unidentified anthocyanins and flavonoids

Anthocyanin glycosylations and acylations create signature fragments upon collision-induced dissociation in the mass spectrometer. Using *m/z* ratios of pelargonidin, cyanidin, peonidin, petunidin, delphinidin and malvidin, and their corresponding glycosylated anthocyanin fragments as baits, we scanned the untargeted LC-MS/MS data from all cultivars for peaks that contained these two fragments at relatively high intensities. This resulted in isolation of 266 peaks, comprising of not just anthocyanins but likely also other flavonoids of similar fragmentation patterns (e.g. glycosides of quercitin, isorhamnetin, chrysoeriol, hesperitin)^13^ **(Supplementary Figure 2)**. From these, we manually identified 67 peaks whose fragmentation patterns were dominated by a single fragment representing the aglycone **(Supplementary Figure 2; Supplementary File 4)** and thus, were likely anthocyanins.

All 266 peaks were clustered using MS/MS molecular networking. Of these, 146 passed the 70% similarity cutoff to generate a network node, with all 18 identified anthocyanin peaks passing all thresholds **(Figure 4E)**. Of the 67 anthocyanin-like peaks, 50 were clustered into 48 nodes in the network. This analysis clearly divided the predicted anthocyanins into sub-networks based on individual aglycone fragments. Specifically, 24, 6, 14, 2 and 2 nodes were associated with the aglycones peonidin, pelargonidin, cyanidin, delphinidin, and petunidin **(Figure 4E)**, respectively, revealing previously uncharacterized diversity in sweet potato anthocyanins. It needs to be noted that the 266 peaks on which this network was based only represent a subset of flavonoids detected in our LC-MS data, given the strict requirement to have a glycosylated fragment of a specific anthocyanidin-associated mass, and thus, several more flavonoids may exist. These results demonstrate the abundance of major and minor flavonoids in purple-fleshed sweet potatoes, reinforcing their status as a health food.

## Discussion

In recent years, better awareness of the nutritional properties of sweet potatoes has increased their popularity among the general public. While the carotenoid, vitamin and mineral content of orange-fleshed sweet potatoes has received more attention, anthocyanins and flavonoids from purple-fleshed varieties also have important health benefits of note^2,12^. Purple-fleshed sweet potatoes are not only attractive for general consumption but are also used in the health food, specialty chemicals, food processing, and cosmetics industries – the latter avenues also possibly fetching higher prices to the growers than traditional grocery markets. It is for these reasons that the cultivation of both orange and purple-fleshed sweet potatoes is being explored in the northern, cooler-climate regions of North America. In this study, we focused on optimizing an extraction protocol and characterizing the anthocyanin content of three purple-fleshed sweet potato varieties that can be grown in these regions.

Comparing these cultivars, we found Kotobuki to have comparable total yield to the orange-fleshed standard Beauregard **(Figure 1A)**, although this was attributed to the plants producing a large number of small-sized, “fingerling” roots, most of which were too small for retail grocery markets. Optimal growing conditions for the northern latitudes will need to be identified for growing such sweet potatoes directly for the consumer markets. In contrast, growing sweet potatoes for anthocyanin extraction is not fettered by their individual size. Our results suggest a yield of ∼390 mg anthocyanins/100g fresh weight of sweet potatoes, which is at the higher end of yields from previous findings **(Table 2)**. We note here that direct comparisons cannot be made between our anthocyanin yields and those from previous studies because of differences in growing/storage conditions, cultivars, and specific extraction steps. Hence, as a rough proxy, we compared across multiple studies to determine where our observed yields lie **(Table 2)**. For example, pressurized liquid extraction using acidified methanol yielded a maximum of 210 mg anthocyanins/100g fresh weight of anthocyanin across 335 sweet potato genotypes grown in North Carolina^48^ and another study found a maximum of ∼96 mg/100g fresh weight across three different cultivars^14^. However, our dry weight average of ∼800 mg anthocyanins/100g of lyophilized powder – which may be a better comparison than fresh weight due to lack of effect of water content – sits in the intermediate to high end of the range of other studies that report values in dry weight for other cultivars^36,37,41,44,48^. This could be because freezing may increase the release of anthocyanins from vacuoles while maintaining their stability and protecting them from degradation by polyphenol oxidase. Lyophilization, however, was found to be a more consistent method than using simply freezing (performed at the same postharvest stage), possibly due to variable water content in the latter. In addition, lyophilization may be a method commercially well-suited for long-term storage and transport. It is also possible that the higher observed yield is influenced by growing in cooler soils – since previous studies in other species suggest this compound class is produced as a response to reactive oxygen species accumulating under cold stress^27–29,49^. However, this hypothesis will need to be formally tested in the future, since cultivar-specific differences and different growth regimens may significantly affect this inference.

**Table 2:** Average anthocyanin yields from lyophilized tissues in this study, compared to previous studies on purple fleshed sweet potatoes using different cultivars.

| Accession name | Yield/100g dry weight | Range of yields/100g dry weights | Yield/100g fresh weight | Range of yields/100g fresh weight |
| --- | --- | --- | --- | --- |
| Purple Passion | 829 | 100-849<sup>41,44,48</sup>, 1592<sup>36</sup>, 1639<sup>37</sup> | 390 | 6.5-210<sup>14,48,50</sup> |
| All Purple Sweet Potato | 776 |  | 381 |  |
| Kotobuki | 765 |  | 399 |  |

In addition to the anthocyanin yield, the types of anthocyanins produced are also important. 16/18 high-confidence anthocyanins we identified across the three purple cultivars were acylated, and >90% of the 67 putative anthocyanins had peonidin or cyanidin as the aglycones. There was no significant difference in anthocyanin concentrations or ratios of cyanidin to peonidin across the tested cultivars, indicating that processes that lead to the accumulation of these compounds are similar. The acylation level of Kotobuki anthocyanins was 30% higher than the other two cultivars, suggesting that this cultivar – also given its other growth characteristics – would be optimal for acylated anthocyanin extraction. Such anthocyanins find use in the food colorant industry, where color stability is important. Purple fleshed sweet potatoes are also rich sources of flavonoids, as evidenced by isolation of >250 flavonoid peaks from LC-MS/MS data **(Figure 4E)**. This result builds upon a previous study, which predicted 56 flavonoids (including 7 anthocyanins) from LC-MS data of different sweet potato cultivars^13^.

In summary, our results suggest that purple sweet potatoes grown in northern latitudes may produce levels of anthocyanins higher or comparable to previous studies performed in different cultivars from warmer regions. However, a formal study comparing growth and nutritional characteristics of the same cultivars grown in warmer and cooler climates is needed. The “fingerling”, slender build of these sweet potatoes may reduce their marketability in grocery stores, however, the high levels of acylated anthocyanins and flavonoids – coupled with their cultivation with reduced or no pesticides – make them attractive for other commercial applications. Further research would be needed to determine optimal cultivars and growing regimens suited for the northern soils and climates.

## Author contributions

Conceived study (GM), designed experiments (AB, GM), performed experiments (AB, KF), analyzed data and wrote the manuscript (AB, GM), provided manuscript feedback (AB, KF, GM).

### Acknowledgements

This work was funded by USDA Hatch grant #1021130 to GM. We are grateful to the Boyce Thompson Institute and Dr. Frank Schroeder for use of the Orbitrap LC-MS, Dr. Mark Sorrells for use of the lyophilizer, and Dr. Jocelyn Rose for use of the spectrophotometer. We especially thank Dr. Steve Reiners and Michael Rosatto for helping us perform field experiments at Cornell AgriTech, Geneva.

## Supplementary Data

**Supplementary File 1:** Phenotyping source data from four sweet potato cultivars.

**Supplementary File 2:** Source data for anthocyanin quantification analysis. Standard curve was generated from serial dilution of cyanin and measured with a spectrophotometer. Molarity (mM) was calculated directly from the linear regression (solving for x) of the standard curve. mg/L was calculated from molarity and used the weight of cyanin as a reference weight. mg total represents the total number of mg in the end 100 or 50 mL extract. This was divided by the number of grams of tissue used in the extraction process to get mg/g concentrations of anthocyanins in sweet potato. Cooking and solvent experiments were all cooked in water and thus fresh weight reporting was not adjusted. Processing methods mixed raw unprocessed samples with samples cooked in water and samples which were lyophilized and lost water and thus have adjustments for water gained and lost.

**Supplementary File 3**: Raw data for anthocyanin quantification from four cultivars. Molarity (mM) was calculated directly from the linear regression (solving for x) of the standard curve. mg/L was calculated from molarity, using the weight of cyanin as a reference weight. mg total represents the total number of mg in the end 100 or 50 mL extract. This was divided by the number of grams of tissue used in the extraction process to get mg/g concentrations of anthocyanins in sweet potato. As all samples were lyophilized, concentrations reflect dry weight and not fresh weight in initial data.

**Supplementary File 4**: MS/MS data of 67 peaks identified as anthocyanin-like compounds, 50 of which are highlighted in **Figure 4E**.

---

**Supplementary Figure 1:**
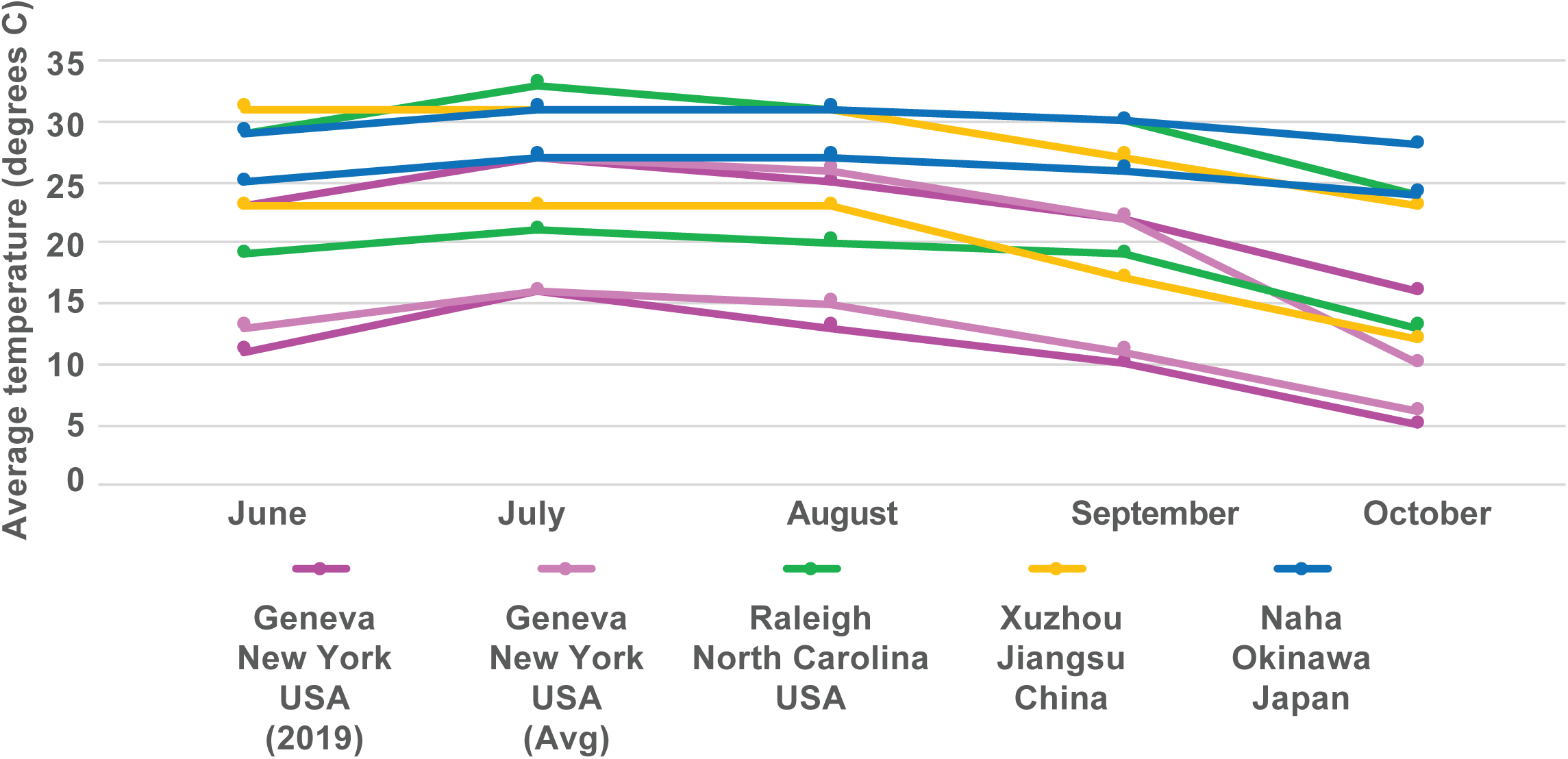
Average high and low temperatures in different sweet potato cultivation areas. The average high and low temperatures at a given location are shown in lines of the same color. Data for Geneva (2019) was obtained from Weather Underground. All other data is high/low averages from 1985-2015 obtained from TimeAndData.com.

**Supplementary Figure 2:**
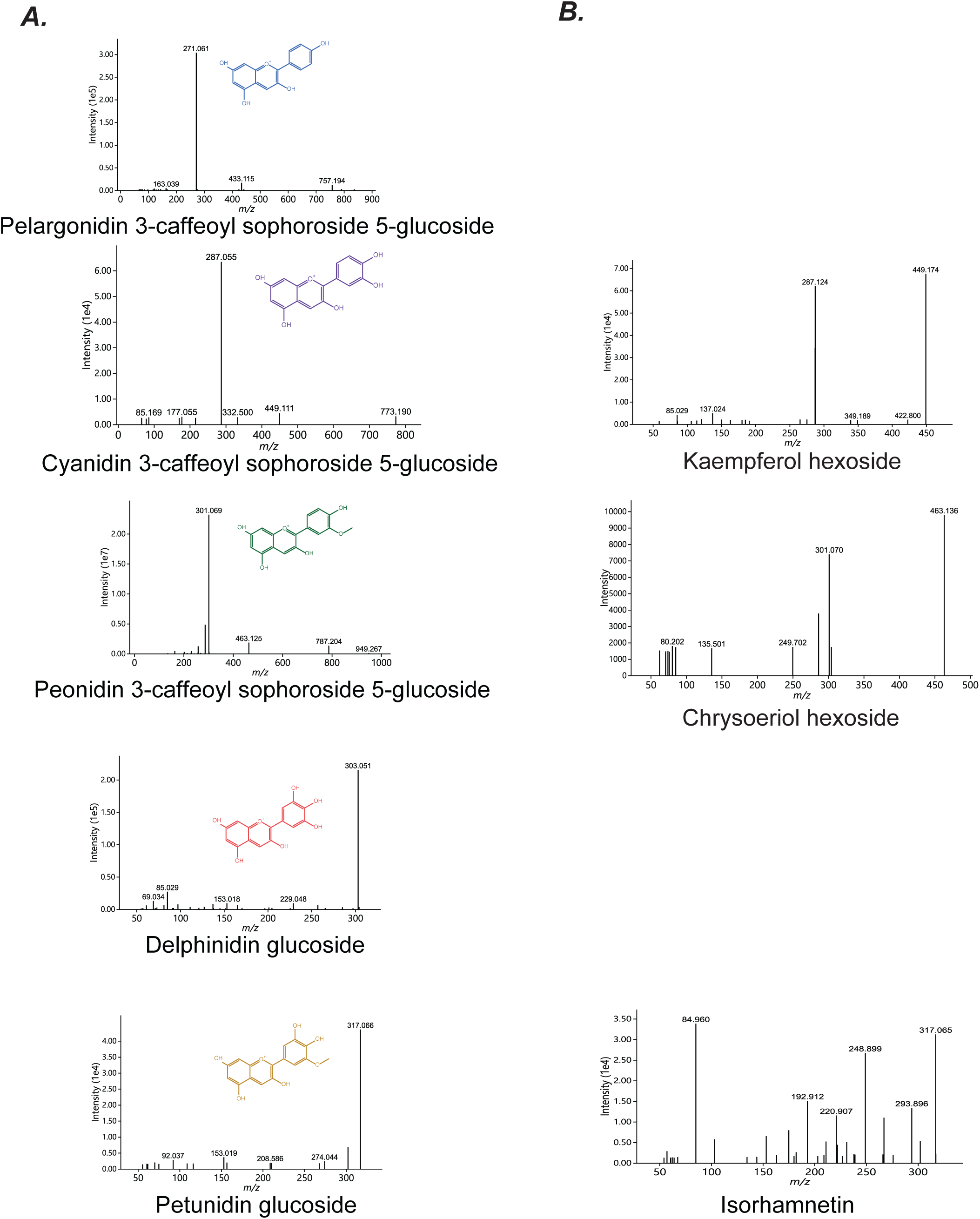
Example MS/MS spectra of (A) anthocyanins and (B) flavonoids of the same parent ion mass detected in our LC-MS datasets and present in the MS/MS molecular network. (A) Spectra of 3-caffeoyl sophoroside 5-glucoside of cyanidin, peonidin, pelargonidin, petunidin, delphidin. The aglycone originating from the flavylium core is the most intense fragment. (B) For the tentatively identified flavonoids, there are typically two intense fragments, one with the glucoside. Identifications are tentative and reflect the most likely flavonoids isomer found in sweet potato.

